# Broadening the scope: Multiple functional connectivity networks underlying threat and safety signaling

**DOI:** 10.1101/2023.08.16.553609

**Authors:** Cody A. Cushing, Yujia Peng, Zachary Anderson, Katherine S. Young, Susan Y. Bookheimer, Richard E. Zinbarg, Robin Nusslock, Michelle G. Craske

## Abstract

**Introduction:** Threat learning and extinction processes are thought to be foundational to anxiety and fear-related disorders. However, the study of these processes in the human brain has largely focused on a priori regions of interest, owing partly to the ease of translating between these regions in human and non-human animals. Moving beyond analyzing focal regions of interest to whole-brain dynamics during threat learning is essential for understanding the neuropathology of fear-related disorders in humans.

**Methods:** 223 participants completed a 2-day Pavlovian threat conditioning paradigm while undergoing fMRI. Participants completed threat acquisition and extinction. Extinction recall was assessed 48 hours later. Using a data-driven group independent component analysis (ICA), we examined large-scale functional connectivity networks during each phase of threat conditioning. Connectivity networks were tested to see how they responded to conditional stimuli during early and late phases of threat acquisition and extinction and during early trials of extinction recall.

**Results:** A network overlapping with the default mode network involving hippocampus, vmPFC, and posterior cingulate was implicated in threat acquisition and extinction. Another network overlapping with the salience network involving dACC, mPFC, and inferior frontal gyrus was implicated in threat acquisition and extinction recall. Other networks overlapping with parts of the salience, somatomotor, visual, and fronto-parietal networks were involved in the acquisition or extinction of learned threat responses.

**Conclusions:** These findings help confirm previous investigations of specific brain regions in a model-free fashion and introduce new findings of spatially independent networks during threat and safety learning. Rather than being a single process in a core network of regions, threat learning involves multiple brain networks operating in parallel coordinating different functions at different timescales. Understanding the nature and interplay of these dynamics will be critical for comprehensive understanding of the multiple processes that may be at play in the neuropathology of anxiety and fear-related disorders.

## Introduction

Responses to threat need to be both fast and accurate to ensure lasting survival in an environment. Learned responses to threat thus need to strike a balance between speed and accuracy. Environmental threats need to be immediately identified to ensure a potentially life-saving reflexive response. However, learning to accurately respond to a threat requires multiple learning episodes to appropriately understand threat contingencies. Deficits in appropriate threat identification or threat response can lead to behavioral outcomes resembling an anxiety disorder (Abend et al., 2022; Britton et al., 2011; Robinson et al., 2013). Consequently, fear-and anxiety-related disorders are often thought to be characterized by aberrations in threat processing and conditioning (Craske et al., 2017; Fenster et al., 2018). For example, anxiety disorders are associated with increased threat acquisition and impoverished threat extinction (Pittig et al., 2018). Anxiety is also thought to be related to how learned threat responses generalize to new stimuli beyond the initial learning episode (Dunsmoor & Paz, 2015). As such, the Pavlovian threat conditioning paradigm has become a pillar in studies examining fear and threat processing due to its simplicity and utility in translational research from animal models to human participants.

Building from animal models, critical brain areas for threat conditioning have been identified such as amygdala and hippocampus (Phillips & LeDoux, 1992). Functioning in these central regions is certainly informative for anxiety disorders as increased anxiety is associated both with facilitated threat responses and difficulty responding to ambiguous threats, as measured from amygdala activation (Im et al., 2017; Pittig et al., 2018). However, important regions like the amygdala are often not detected as having a role in human fMRI studies of threat conditioning (Fullana et al., 2016; Visser et al., 2021; Young et al., 2021). Consequently, it is necessary to expand beyond these critical nodes to the larger functional networks in which they operate.

Especially at the human level, it is likely that there are multiple circuits operating in parallel in what we would typically label the “fear” response (LeDoux & Pine, 2016). We use the term “threat conditioning” here - while noting that much of the literature traditionally refers to this paradigm as “fear conditioning” - as a means of capturing the multidimensional response to learned threat, which includes the subjective experience of fear as well as other physiological and behavioral responses (Mobbs et al., 2009; Taschereau-Dumouchel, Michel, et al., 2022). Multiple circuits underlie the complex of physiological, behavioral and subjective responses and their coordination for an adaptive or coherent response suitable for level of threat imminence (Bolles & Fanselow, 1980; Mobbs et al., 2007). As these processes may potentially become dissociated during certain treatments - e.g. amygdala and skin conductance response reduction with no corresponding reduction in self-reported fear (Cushing, Lau, et al., 2023; Taschereau-Dumouchel et al., 2018) – it is important to understand the neural circuitry behind each of these processes to most effectively tailor future treatments (Cushing, Dawes, et al., 2023; Taschereau-Dumouchel, Cushing, et al., 2022; Varkevisser et al., 2023).

Much of the previous human neuroimaging work using the Pavlovian threat conditioning paradigm has used univariate contrasts to identify threat-sensitive brain regions (Fullana et al., 2016). However, whole-brain connectivity is beginning to be used to understand the broader large-scale dynamics at play in human threat conditioning (Berg et al., 2021; Wen et al., 2021). Network analyses have become increasingly popular as a way to understand how distributed regions across the entire brain organize their activity in coordinated functions (Sporns, 2014). Understanding how large-scale brain networks operate during threat conditioning is likely to advance our understanding of the psychopathology of anxiety and fear-based disorders (Bressler & Menon, 2010; Menon, 2011). By understanding these large-scale dynamics, more effective neurobiologically-targeted treatments may be developed by seeking to stabilize brain activity in these networks as a whole rather than trying to change complex symptomology by affecting singular brain regions in isolation.

Salience and central executive networks have been shown to track overgeneralization of conditioned fear responses in post-traumatic stress disorder (Berg et al., 2021). Extinction of conditioned threat has also been shown to modulate brain connectivity in areas associated with default mode, frontoparietal, and ventral attention networks (Wen et al., 2021). These recent endeavors represent some of the first attempts to look outside the canonical defensive threat/fear network regions to understand threat conditioning processes at a holistic whole-brain scale. Additionally, by examining connectivity these recent studies are moving beyond just understanding which brain regions are involved in threat conditioning. They are exploring the important question of how brain regions coordinate with each other to form new memories about threat and fear.

Here, we add to this growing body of research by utilizing data-driven group independent component analysis (ICA) methods to investigate functional connectivity in brain networks involved in the acquisition and extinction of conditioned threat, as well as the recall of extinction memory after a 48-hour consolidation period. Group ICA has been established as a method for determining functional connectivity patterns in fMRI without having to rely on specific models of brain activity in both resting state and task data (Calhoun et al., 2001, 2009; Igelstrom et al., 2015; Webb et al., 2016). By analyzing over 200 participants, this study represents one of the largest and most robust investigations of whole-brain dynamics during threat conditioning. Moreover, using group ICA, we are able to categorize multiple distinct, data-driven connectivity networks operating in parallel allowing us to characterize different threat and safety sensitivities during threat conditioning.

Such methodology allows us to expand upon and provide additional insight into other recent explorations in whole-brain connectivity during threat conditioning, such as one that explored the temporal dynamics of fear learning and extinction at the single-trial level (Wen et al., 2021). This study was limited in its ability to isolate specific functional networks due to the nature of ROI-to-ROI pair-wise connectivity measures. The group ICA approach presented herein supplements Wen et al. (2021) by dividing whole-brain connectivity into spatially independent, temporally parallel networks and examining functional connectivity during threat conditioning within each one. Additionally, instead of relying upon models of pre-defined regions or anatomically restricted parcellations, the data-driven nature of group ICA allows the data itself to determine which groups of brain regions are working in concert during threat and safety learning.

We predict we will find multiple task-related functional connectivity networks during our 2-day threat conditioning task with this strategy. We expect to largely corroborate the findings of other previous large-scale connectivity studies with independent connectivity networks overlapping or resembling the default mode, salience, and frontoparietal control networks (Berg et al., 2021; Marstaller et al., 2017; Wen et al., 2021) while also potentially finding novel results in additional networks due to the model-free nature of our planned analysis. Importantly, our use of group ICA on the threat conditioning task data itself (as opposed to resting state data) will allow us to examine specifically which portions of these previously implicated canonical networks are engaged during the threat conditioning task. In addition, it enables us to assess conditioned stimulus (CS) response profiles in these functional connectivity networks in a clear-cut network-specific fashion as each connectivity network is defined by a singular timecourse of BOLD activity that is relatively free from the influence of motion or other noise sources.

## Methods

### Participants

Participants were enrolled as part of a cross-site Brain, Motivation, and Personality Development (BrainMAPD) study at University of California, Los Angeles and Northwestern University that has been described in previous research (Anderson et al., 2023; Peng et al., 2023; Rosenberg et al., 2021; Young et al., 2021). We recruited 272 participants (mean age (s.d.) = 19.16 (0.52), 182 female). Participant recruitment focused on participants 18-19 years old and was structured to ensure an even distribution across threat and reward sensitivity, measured by the Behavioral Activation Scale (BAS) and Eysenck Personality Questionnaire-Neuroticism (EPQ-R-N) (Carver & White, 1994; Eysenck & Eysenck, 1993). Participants were ineligible if meeting any of the following criteria: not right handed, not fluent in English, history of traumatic brain injury, MRI contraindications, pregnancy, color blindness, lifetime psychotic symptoms, bipolar I disorder, substance use disorder in the past 6 months, or currently taking antipsychotic medication. Participants provided informed consent according to the procedures approved by UCLA and NU Institutional Review Boards. Participants were excluded from all fMRI analysis if they demonstrated excessive motion (defined as >10% outlier scans) in any of the 3 task phases (acquisition, extinction, and extinction recall). After exclusions for motion and technical issues during data collection, 223 participants were analyzed for acquisition and 208 participants were analyzed for extinction and extinction recall.

### Task

Participants completed a 2-day, three-phase (acquisition, extinction, and extinction recall) Pavlovian threat conditioning task as described previously (Peng et al., 2023; Rosenberg et al., 2021; Young et al., 2021) while undergoing an fMRI scan. Acquisition and extinction occurred sequentially during the same scanning session while extinction recall was conducted in a separate fMRI session at least 48 hours later. Each trial began with 3 seconds of a context image presentation (office or conference room setting with non-colored lamp) followed by 6 seconds of CS presentation (lamp color changing to red, yellow, or blue). During acquisition, the unconditioned stimulus (US) was delivered via electric shock that co-terminated with the CS. Stimulus offset was followed by a varied inter-trial interval of 12 to 18 seconds (mean 15 seconds). Acquisition occurred in one context while extinction and extinction recall occurred in the other context. Context and CS lamp-color assignments were counterbalanced across participants. Acquisition consisted of 8 trials of a CS+ that became the extinguished CS (CS+E), 8 trials of a CS+ that would remain unextinguished (CS+U) and a CS-that was never paired with shock. CS+ stimuli were reinforced with shock on 5 out of 8 trials each. Extinction contained 16 CS+E trials and 16 CS-trials. Extinction recall contained 8 CS+E trials, 8 CS+U trials, and 16 CS-trials. No shocks were delivered in either the extinction or extinction recall phases.

### MRI acquisition parameters

MRI data were collected using a PRISMA 3T scanner with a 64-channel head coil in a cross-site study at the UCLA Ahmanson-Lovelace Brain Mapping Center and the Northwestern University Center for Translational Imaging. Acquisition parameters were identical at both sites. High resolution T1-weighted structural images were collected using a magnetized prepared rapid acquisition gradient echo (MPRAGE) sequence (voxel size=0.8 mm^3^, TR/TE/flip angle=2300ms/2.99ms/7°, FOV=256mm^2^, 208 slices). Functional images were acquired parallel to the AC-PC line with a multiband sequence (voxel size=2.0mm^3^, TR/TE/flip angle=2000ms/25ms/80°, FOV=208mm^2^, 64 slices, multiband acceleration factor=2) collecting 380 volumes for each task phase.

### MRI preprocessing

Structural T1 images were intensity normalized and the brain extracted using optiBET (Lutkenhoff et al., 2014). The brain extracted T1 image was then segmented into White Matter, Gray Matter, and Cerebrospinal Fluid using FAST (Y. Zhang et al., 2001).

Functional images were preprocessed separately for the acquisition, extinction, and extinction recall runs. Motion outliers were calculated for functional images using *fsl_motion_outliers* before any preprocessing took place. Then, functional images were motion corrected, smoothed with a 4 mm FWHM kernel, and nonlinearly registered into the MNI152NLin6Asym standard space using 12 degrees of freedom with FSL’s FEAT (FMRIB’s Software Library, www.fmrib.ox.ac.uk/fsl). Motion components were then automatically detected and removed using ICA-Aroma (Pruim et al., 2015). ICA-aroma was used as it has been found to be one of the most effective methods of removing motion from fMRI data without discarding large amounts of data (Parkes et al., 2018). Following ICA-aroma, data had linear and quadratic trends removed along with 6 head motion parameters from FSL and white matter and cerebrospinal fluid time courses using AFNI (Cox, 1996; Cox & Hyde, 1997). A highpass filter was also applied to remove frequencies below .008 Hz during this step to remove potential sources of noise. A low-pass filter was not applied to prevent the filtering of any task-relevant content contained in the high frequencies. Finally functional data were transformed into the standard space using parameters from the FSL non-linear registration.

### Group ICA connectivity and dual regression

Similar to preprocessing, a separate group ICA connectivity analysis was performed for the acquisition, extinction, and extinction recall fMRI runs. Preprocessed functional data for all participants were submitted to group ICA in Melodic from FSL using the temporal concatenation method. Group ICAs were limited to 20 ICs based on previous work (Webb et al., 2016) and to limit the number of multiple comparisons. The group ICA analysis was masked to brain tissue only using the FSL standard brain mask. Group IC maps were thresholded at Z=4 for generation of figures and identification of key network regions. Any components that resembled physiological noise or did not have significant contributions from the cortex (e.g. cerebellar networks) were discarded from further analysis. This resulted in 16 connectivity networks analyzed during each task phase. Dual regression was also performed using FSL to obtain participant-specific time course contributions to each IC.

### Modeling IC connectivity response to task conditions

To find how connectivity within each IC responded to the conditions of the task, a GLM was fitted to each participant-specific time course for each IC using pyMVPA in python (Hanke et al., 2009). This process is identical to modeling a typical univariate whole-brain GLM but rather than a voxel time course being used as a dependent variable, the time course of an IC is used. Specifically, the conditions of CS+ and CS-were modeled separately for the early and late phases of each run for acquisition and extinction. The unconditioned stimulus (US) was also modeled during acquisition. For extinction recall, early and late unextinguished CS+ (CSU), extinguished CS+ (CSE), and CS-were modeled. Early and late represented the first and last 4 trials of each stimulus type, respectively. A regressor was also included for context presentation (office or conference room image) during each experimental phase. Motion outliers identified in preprocessing were included as additional nuisance regressors in the GLM to minimize effects of movement on parameter estimation. GLMs were modeled with an SPM-style hemodynamic response function model including a temporal derivative to account for temporal differences in slice acquisition. All regressors were specified with a duration of 0 seconds at the onset of each stimulus for an event-related response over epoch response in order to minimize influence of temporally adjacent events.

### Statistical Analysis

To identify which IC networks had functional connectivity that was sensitive to task conditions, a mixed linear model was calculated for each IC within each task using the statsmodels package in python. In acquisition and extinction, the dependent variable of IC Beta estimate was tested for the interaction of fixed effects condition (CS+/CS-) and time (early/late) for each IC. For extinction recall, we were only interested in the early phase of the task so a model of fixed effect of condition (early CSU vs. early CSE) was tested. Participant and scan site (UCLA/NU) were included as random effects with participant nested within scan site for each model. With a model for each IC not discarded for resembling noise/non-cortical sources, 16 total models were tested for each task phase. Results of each model were Bonferroni corrected for multiple comparisons.

### Canonical Network Overlap

In order to characterize with which canonical networks each IC network identified in our analyses overlapped, we calculated the overlap between our IC spatial maps and 7 canonical networks (Schaefer et al., 2018): Visual, Dorsal Attention, Control, Default Mode, Limbic, Salience/Ventral Attention, and Somatomotor. Canonical network masks were generated by combining and binarizing individual parcels belonging to each of the 7 networks from the 100 parcel 7 network Schaefer atlas (Schaefer et al., 2018). Thresholded IC spatial maps (described above) were binarized. Overlapping voxels between the thresholded IC maps and each network mask were determined by multiplying the 2 binarized images together. Then, the percentage of overlap between each IC network and each canonical network was calculated by dividing the number of overlapping voxels by the total number of voxels in each canonical network mask.

## Results

### Threat Acquisition

During threat acquisition there was a significant interaction between CS-type and Time (*F*(1,888)=15.89, *p*=0.0012 Bonferroni corrected) in a network, referred to as network 1, including vmPFC, OFC, hippocampus, angular gyrus, posterior cingulate, and retrosplenial cortex overlapping with the Default Mode Network (Fig. 1A). This interaction was characterized by no initial difference in connectivity between CS+ and CS-during early trials (*t*(222)=0.99, *p*=0.32) while in late trials network connectivity was significantly greater for CS-compared to CS+ (*t*(222)=6.71, *p*<0.001). Such late decreases in response to CS+ are thought to identify regions important for threat extinction (Garcia et al., 1999; Hennings et al., 2020; Phelps et al., 2004).

**Figure 1.**
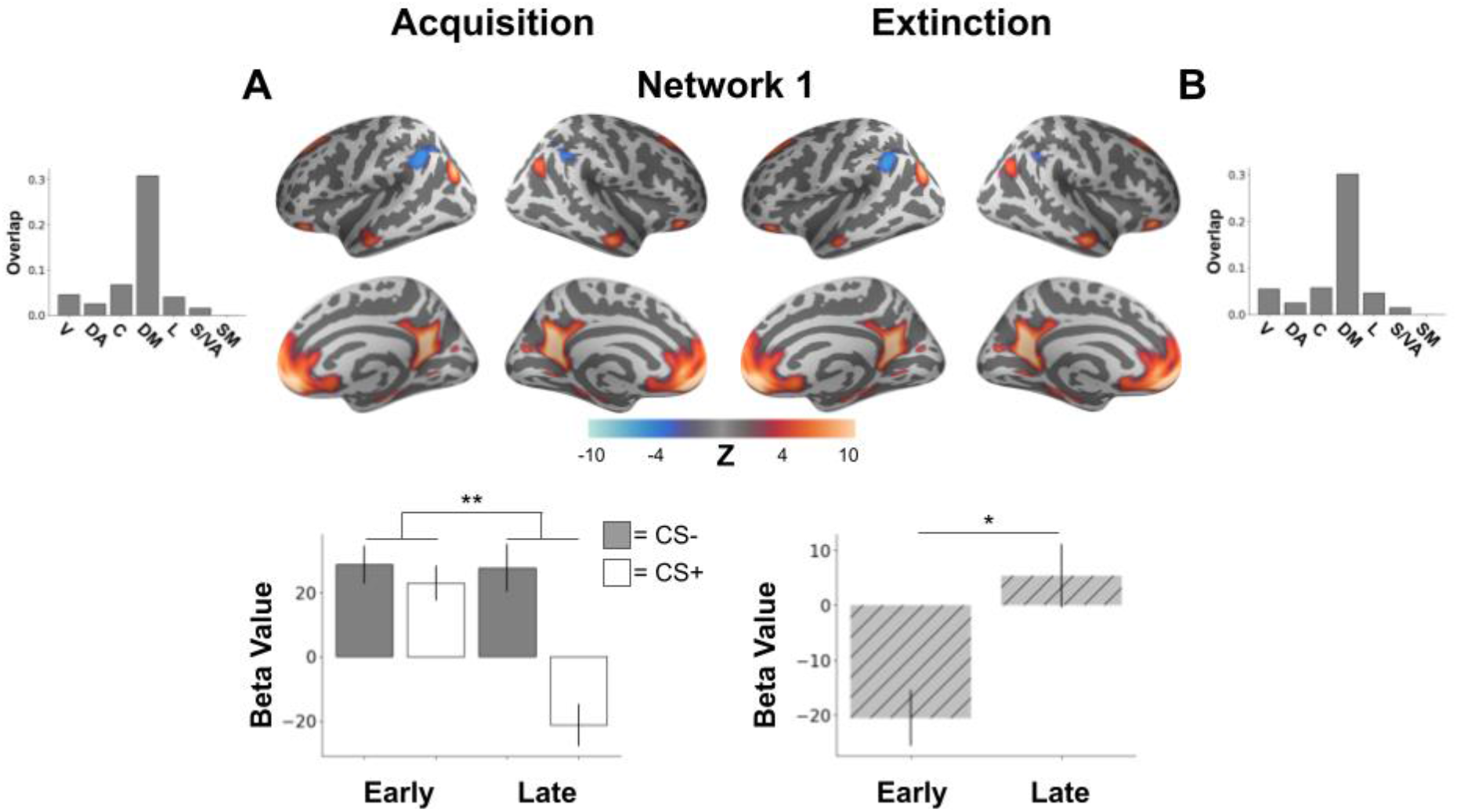
Brain network demonstrating acquisition and extinction of learned threat. Brain plots show thresholded independent component (IC) spatial maps. Bottom bar plots show IC-specific GLM parameter estimates. Top bar plots show connectivity network overlap with canonical networks. (A) A distributed brain network involving bilateral hippocampus, vmPFC, and posterior cingulate demonstrated a significant interaction between CS-type (CS-/CS+) and time (early/late) during threat acquisition. During late acquisition, the network demonstrated increased connectivity to the CS-compared to the CS+. (B) This same brain network involving bilateral hippocampus, vmPFC, and posterior cingulate was observed during threat extinction with connectivity in the network increasing from early to late extinction. * *p*<0.05, ** *p*<0.01, V=Visual, DA=Dorsal Attention, C=Control, DM=Default Mode, L=Limbic, S/VA=Salience/Ventral Attention, SM=Somatomotor

Additionally, two networks exhibited a main effect of CS-type. These networks demonstrated the canonical threat acquisition response of CS+>CS-indicating successful acquisition of threat-related response. A network, designated network 2, included dorsal anterior cingulate cortex (dACC), mPFC, and inferior frontal gyrus (*F*(1,888)=15.70, *p*=0.001 Bonferroni corrected, Fig. 2A). The other network, network 4, spanned the entirety of insular cortex including anterior insula along with cingulate gyrus, inferior frontal gyrus and OFC (*F*(1,888)=109.91, *p*<0.001 Bonferroni corrected, Fig. 3B). Both networks showing threat sensitivity overlapped most with the salience/ventral attention network, indicating the salience network’s involvement in learned threat detection.

**Figure 2.**
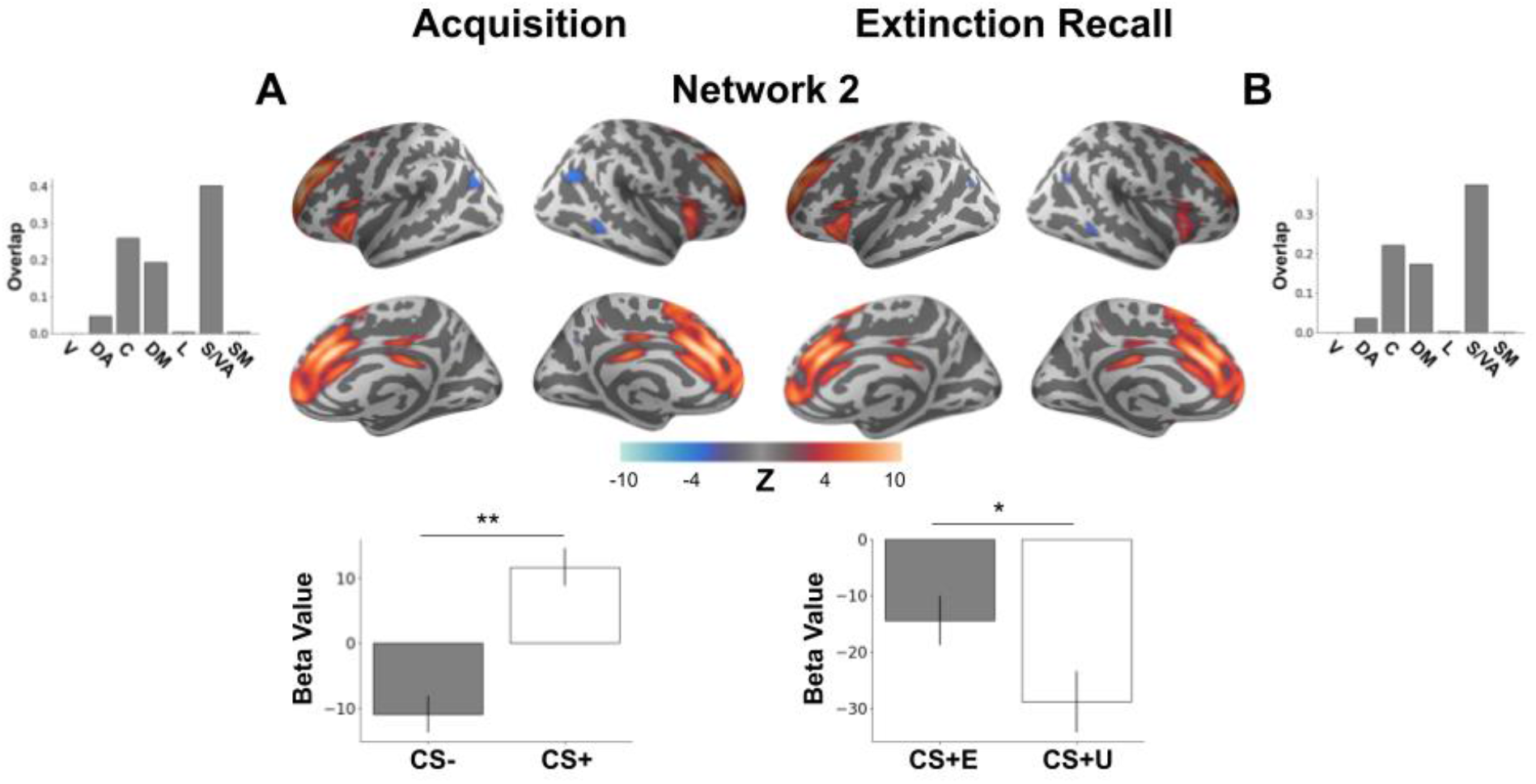
Brain network demonstrating acquisition of learned threat and recall of extinction memory. Brain plots show thresholded independent component (IC) spatial maps. Bottom bar plots show IC-specific GLM parameter estimates. (A) A distributed brain network consisting of dorsal anterior cingulate cortex (dACC), mPFC, and inferior frontal gyrus demonstrated an effect of CS-type during threat acquisition with greater connectivity elicited by the CS+ compared to the CS-. (B) This same brain network was observed during extinction recall with significantly decreased connectivity elicited by the unextinguished CS+ (CS+U) compared to the extinguished CS+ (CS+E). * *p*<0.05, ** *p*<0.01, V=Visual, DA=Dorsal Attention, C=Control, DM=Default Mode, L=Limbic, S/VA=Salience/Ventral Attention, SM=Somatomotor

**Figure 3.**
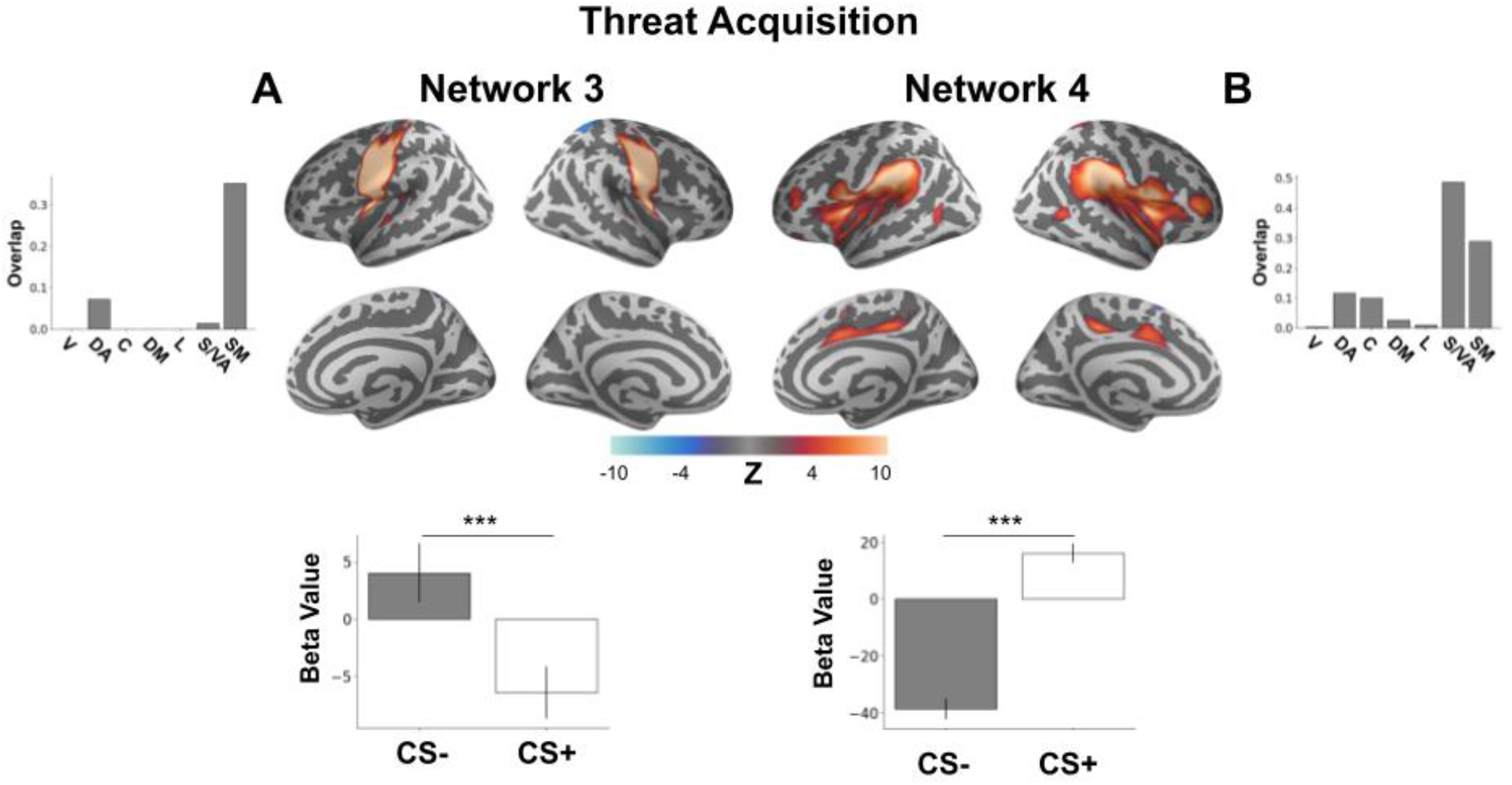
Brain networks involved in learned threat acquisition. Brain plots show thresholded independent component (IC) spatial maps. Bottom bar plots show IC-specific GLM parameter estimates. Top bar plots show connectivity network overlap with canonical networks. (A) A brain network involving precentral and postcentral gyri as well as the left insular cortex demonstrated increased connectivity elicited by the CS-and decreased connectivity elicited by the CS+ during threat acquisition. (B) A brain network involving the insula, middle frontal gyrus, cingulate gyrus, and OFC. This network demonstrated acquired threat response with increased connectivity induced by the CS+ and decreased connectivity elicited by the CS-. *** *p*<0.001, V=Visual, DA=Dorsal Attention, C=Control, DM=Default Mode, L=Limbic, S/VA=Salience/Ventral Attention, SM=Somatomotor

Another network, network 3, demonstrated a main effect of CS-type with a similar pattern of CS-connectivity being significantly greater than CS+ connectivity as observed in Fig. 1A (*F*(1,888)=14.13, *p*<0.001 Bonferroni corrected). The network consisted of the precentral and postcentral gyri and insular cortex, overlapping with the Somatomotor network (Fig. 3A).

### Threat Extinction

Interestingly, there was no CS-type sensitivity during threat extinction despite immediately following threat acquisition. No IC networks demonstrated interactions of CS-type and time or main effects of CS-type. This may due to a dishabituation effect for the CS-due to the break between fmri scans following acquisition and before extinction. However, extinction processes could be tracked through observed main effects of time.

Most saliently, we observed a main effect of time in the same network 1 involving vmPFC, hippocampus, and posterior cingulate from threat acquisition (Fig. 1A) during extinction (*F*(1,828)=11.42, *p*=0.012 Bonferroni corrected, Fig. 1B). Network 1, overlapping with Default Mode Network demonstrated decreased connectivity in response to CS stimuli during early extinction that increased to positive connectivity during late extinction. This change in connectivity likely tracked extinction learning due to the positive increase over the extinction period as opposed to a decrease over the extinction period as might be expected from habituation or unrelated processes. Another network, network 5, overlapping with the visual network involving lateral occipital cortex and fusiform cortex showed the same connectivity increase from negative to positive over the extinction period (*F*(1,828)=30.04, *p*<0.001 Bonferroni corrected, Fig. 4A). Though this network was not observed during the threat acquisition process, this network activity also potentially tracks extinction learning due to the increase of connectivity over the task period.

**Figure 4.**
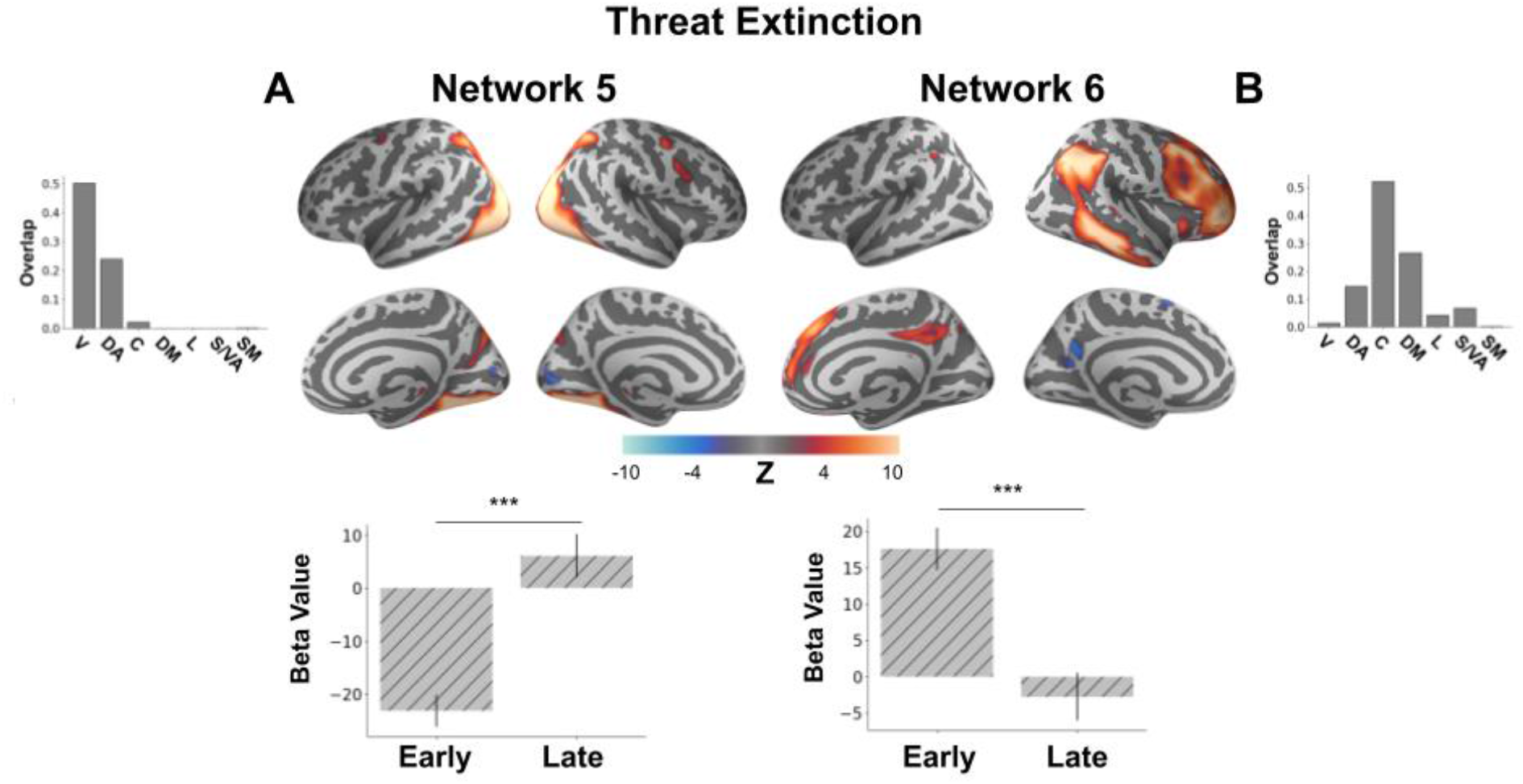
Brain networks involved in the extinction of learned threat response. Brain plots show thresholded independent component (IC) spatial maps. Bottom bar plots show IC-specific GLM parameter estimates. Top bar plots show connectivity network overlap with canonical networks. (A) A brain network involving lateral occipital cortex as well as fusiform cortex demonstrates a significant increase in connectivity in response to CS stimuli from early to late trials in the extinction learning phase. (B) A brain network involving angular gyrus, frontal gyrus, and posterior cingulate showed significantly reduced connectivity in response to CS stimuli from early to late trials in the extinction learning phase. ***p<0.001, V=Visual, DA=Dorsal Attention, C=Control, DM=Default Mode, L=Limbic, S/VA=Salience/Ventral Attention, SM=Somatomotor

Finally, we observed a main effect of time in another network, network 6, involving angular gyrus, frontal gyrus, and posterior cingulate, overlapping most with the Control network (*F*(1,828)=21.49, *p*<0.001 Bonferroni corrected, Fig. 4B). However, as the connectivity pattern of this network decreased from early extinction to late extinction, it is difficult to attribute this result to extinction learning directly without stimulus-specific evidence as it could also represent stimulus habituation or other unrelated processes.

### Extinction Recall

As only the early period of extinction recall was of experimental interest, we ran a model examining only the main effect of CS-type in the early phase of the task (early CS+E vs. early CS+U). This revealed the same network found during threat acquisition, network 2, which exhibited a greater decrease in connectivity (*F*(1,412)=10.15, *p*=0.025 Bonferroni corrected) for the unextinguished CS+ (CS+U) compared to the extinguished CS+ (CS+E, Fig. 2B). These results showcase the importance of this network both in the acquisition of threat learning as well as the expression of extinction memory.

## Discussion

In this study, we examined functional connectivity networks during a 2-day Pavlovian threat conditioning paradigm using fMRI. With a data-driven group independent component analysis, we compared how spatially independent functional connectivity networks responded to conditioned stimuli during acquisition and extinction of threat responses as well as extinction recall a full 48 hours later. This revealed multiple distinct brain networks involved in the acquisition, extinction, and recall of extinction memory for learned threat. A stable network, network 1, overlapping with the Default Mode Network involving hippocampus, vmPFC, and posterior cingulate was involved in both the acquisition and extinction of learned threat. An additional persisting network, network 2, overlapping with the Salience Network involving dACC, mPFC, and inferior frontal gyrus was involved in the acquisition of learned threat as well as the expression of extinction memory during extinction recall. Additional networks overlapping with parts of the salience/ventral-attention and somatomotor networks were involved in the acquisition of learned threat responses. Portions of the visual and fronto-parietal control networks were implicated in threat extinction.

The finding of network 1, involving hippocampus and vmPFC, is consistent with previous work that has focused on these regions as part of a ‘network’ that consistently responds to threat conditioning paradigms (Giustino & Maren, 2015; Picó-Pérez et al., 2019). It is worth noting that these regions were found to be functionally connected in the current work without the imposition of a model through the a priori selection of regions of interest. This strengthens the argument for them as canonical threat learning regions while demonstrating the coordination of these regions in a self-contained functional connectivity network. Responses specific to the conditioned cue developed late in the threat acquisition phase, indicating network 1’s involvement in the learning aspect of threat acquisition. Connectivity in this network then increased during the extinction process, again suggesting a learning process over the extinction period. As the observed network partially overlaps with the canonical default mode network, this adds to a building body of evidence in human neuroimaging implicating the default mode network in threat learning (Berg et al., 2021; Marstaller et al., 2017; Wen et al., 2021; Zidda et al., 2018). Additionally, this supports the default mode network as a potential hub in the pathophysiology of anxiety and fear-based disorders (Marstaller et al., 2017; Menon, 2011; Son et al., 2023; J. Zhang et al., 2023; Zidda et al., 2018). As this network was involved in threat learning and extinction in the current study, it is a prime candidate for future studies concerning the relation of this network to the pathophysiology of anxiety disorders and fear-related disorders.

We also observed a stable network (network 2) across threat acquisition and extinction recall in dACC, inferior frontal gyrus, and mPFC overlapping partly with the salience network. As the observed stimulus specificity during acquisition in this network was not specific to the late acquisition period, it is possible that this network detects learned threats at a rapid rate (e.g. one-shot learning). This network was also the only network to show conditioned stimulus specificity (between the extinguished and unextinguished conditioned stimuli) following the acquisition period. As this stimulus specificity was observed 48 hours after acquisition and extinction, it is implicated in long-term storage of learned threat responses. This underscores the importance of this network of regions in the threat learning process as a site of potentially both rapid and lasting threat memory acquisition. These results may be of broad clinical significance in understanding anxiety and fear-related disorders as the salience network itself has been found to track fear generalization and symptom severity in post-traumatic stress disorder (Berg et al., 2021). Perhaps a rapid learning rate within this network predisposes it to overgeneralized and persistent threat responses.

Rapid threat learning within the salience network is further supported by the current finding of an additional insula-centered network (network 4) during threat acquisition where conditioned stimulus specificity emerged early. The salience network involving the insula is thought to be responsible for detecting salient events in a bottom-up fashion while coordinating other networks to access attention, working memory, and motor systems in response to salient events (Uddin, 2015). Consequently, the salience network demonstrating threat learning early in the acquisition phase supports this model of function, as the CS+ stimulus should be salient following the first delivery of the US. As functional connectivity was greater for the CS+ compared to the CS-in both of our salience/ventral attention-overlapping networks during acquisition, this may indicate greater bottom-up processing for the CS+ in these networks.

In addition, a functional connectivity network overlapping primarily with the somatomotor network (network 3) involving pre-and post-central gyri was involved in threat acquisition. As these regions correspond to the motor and somatosensory cortices, connectivity in this network may reflect the preparation of motor responses to threat. Important to note, functional connectivity in this network was greater for the safety stimulus, CS-, compared to the threat stimuli, CS+. A recent meta-analysis similarly found greater activation in the primary somatosensory cortex to the CS-versus the CS+ (Fullana et al., 2016). So, this finding may conversely represent a safety signal, relating to approach-oriented motor behaviors.

During threat extinction, beyond the network overlapping with the default mode network, we observed two functional connectivity networks overlapping with the visual (network 5) and fronto-parietal control (network 6) networks. Functional connectivity in the visual network involving lateral occipital and fusiform cortices demonstrated an increase in connectivity during extinction from early to late trials. This increase in functional connectivity may represent processing of visual contextual information to help process the new safe context as well as the CS+’s transition itself to a “safe” stimulus. Conversely, functional connectivity in this fronto-parietal control network decreased from early to late trials during extinction. As the fronto-parietal network is associated with top-down regulation and task flexibility (Cole et al., 2013; Dosenbach et al., 2006; Power et al., 2011; Rossi et al., 2009), this decrease in connectivity may represent a reduction in top-down regulation or attentional processes that are necessary during early extinction when it is uncertain for participants whether stimuli may still represent a threat. However, as this was a general decrease in connectivity across both stimulus types, this decrease in connectivity may simply reflect stimulus habituation or other unrelated processes.

Interestingly, we did not observe conditioned stimulus specificity during the extinction period in any examined network. This is all the more perplexing given the extinction phase immediately followed the threat acquisition phase, in which conditioned stimulus specificity was widely observed. Nonetheless, the result broadly matches other whole-brain connectivity findings using a similar paradigm (Wen et al., 2021) in which no conditioned stimulus-specific response was observed early in extinction (though, unlike our study, they did observe stimulus specificity late in extinction). One potential explanation is a dishabituation effect for the CS-due to the break between fMRI scans following acquisition and before extinction. Other studies have observed a heightened sensitivity to the CS-stimulus at the beginning of extinction when there is an interruption between acquisition and extinction (Zbozinek et al., 2022). Another possible explanation relies on contextual effects, since extinction always took place in a different context than acquisition. The change in context between acquisition and extinction may have overshadowed the discrete stimuli (Fanselow, 2010). However, as there was conditioned stimulus specificity during extinction recall in the same extinction context 48 hours later, context presentation alone does not sufficiently explain the lack of stimulus specificity in extinction. Another potential explanation is broad stimulus habituation effects due to the close temporal proximity of acquisition and extinction, resulting in loss of specificity by the time of extinction. Moreover, consolidation processes that enable stimulus specificity may have required the 48 hours before recall, resulting in no specificity at the end of extinction but specificity at the beginning of recall.

Strengths of our study include a large sample size of more than 200 participants. As neuroimaging frequently suffers from low sensitivity and under-powered sample sizes (Thirion et al., 2007), the findings here can be considered robust. Despite this high-powered sample, significant amygdala involvement was conspicuously missing from any of the networks found in this current analysis. This matches large meta-analyses as well as other findings that find a minimal role (if any) for the amygdala in human fear conditioning paradigms (Fullana et al., 2016; Visser et al., 2021; Young et al., 2021). The lack of significant activity in the amygdala during threat conditioning paradigms in human neuroimaging has been attributed to less salient threat in human studies compared to animal studies, as well as a general difficulty in measuring transient responses with fMRI (Fullana et al., 2016, 2019; Somerville et al., 2013). Moreover, as we have argued elsewhere, emotions evoked by threat versus safety may rely upon distributed networks rather than single regions such as the amygdala. On the other hand, with enough statistical power, reliable contributions to threat conditioning and safety learning can be observed in the human amygdala with fMRI (Wen et al., 2022).

The current study is not without limitations, however. We recognize that the totality of these results across many networks do not necessarily congeal into a singular cohesive narrative. By using a data-driven method like group ICA with minimal network inclusion criteria, we are reporting results that may not immediately fit into canonical theories or thinking about threat conditioning. As a result, the findings independently validate the recent ideas that the default mode and salience networks are important for human threat conditioning in a model-free fashion (Berg et al., 2021; Marstaller et al., 2017, 2021). Consequently, this collection of results may prove important for expanding the field beyond the traditional fear network regions, pending further replications and extensions relating brain networks to behavioral responses and symptom dimensions.

Additionally, this cross-sectional dataset is limited in the conclusions that can be drawn. Participants analyzed in this study were relatively young (average age of 19) with low variance in age due to this data coming from a larger, longitudinal study looking at developmental changes from adolescence into early adulthood. In the future, a longitudinal analysis of functional connectivity during threat learning and extinction will inform the stability of these connectivity networks over time and how their function relates to development of symptoms around anxiety and fear-based disorders. The results of this study are also limited in what they can say about the subjective experience of fear itself, as participants were not asked to rate their fear levels during or after the task. Consequently, the results of this study should be interpreted in the lens of general threat responses encompassing both non-conscious physiological threat responses and consciously experienced fear and threat awareness. Additionally, despite our large sample size, our data-driven group ICA analyses were heavily corrected for multiple comparisons. While this ensures the results reported here are robust, subtle effects may have been mitigated, which may explain why the amygdala was not detected in our functional connectivity networks (Wen et al., 2022) or why stimulus specificity was not detected during threat extinction despite other recent results showing stimulus-specific connectivity increases at the end of extinction (Wen et al., 2021).

In summary, using data-driven methods we have characterized several functional connectivity networks involved in the Pavlovian threat conditioning paradigm. These functional connectivity networks overlap most commonly with areas involved in the default mode and salience networks, highlighting the potential importance of these canonical networks in acquisition, extinction, and extinction recall processes. This supports other recent findings from whole-brain connectivity analyses of fear conditioning that demonstrate the need for researchers to move beyond the previously focal region of interest analyses of fear conditioning paradigms in order to understand the dynamic nature of the human brain’s response in threat learning (Wen et al., 2021). The present work has the added benefit of refining these whole-brain functional connectivity patterns into spatially independent networks to identify which regions work directly in concert as well as which functional connectivity networks are involved in different aspects of threat learning and extinction. Future work will need to disentangle which of these networks are involved in automatic defensive responses to threat and which contribute to the actual subjective experience of fear and threat in the human brain in order to better tailor clinical interventions for fear-related disorders to meet individual needs or symptom profiles.

